# Path analysis reveals that corticosterone mediates gluconeogenesis from fat-derived substrates during acute stress in songbirds

**DOI:** 10.1101/2023.01.24.525409

**Authors:** H. Bobby Fokidis

## Abstract

Glucocorticoids (e.g., corticosterone or CORT in birds) mobilize energy reserves during stress to aid survival. Stress liberates glucose (GLU) by glycogenolysis, but with glycogen depletion, gluconeogenesis of fat and protein sources predominates. Songbirds have higher metabolic rates and GLU concentrations than mammals and likely rely more on fat and protein stores during stress. We tested this hypothesis in four songbird species using path analysis to model the interrelationships between CORT and energy metabolites both at baseline and after acute stress. Individuals in better condition had higher triglyceride and CORT levels at baseline than individuals in poor body condition, and these differences became more pronounced with stress. Free CORT (the fraction unbound to circulating proteins) was associated with more GLU and free glycerol at baseline, but the former relationship was lost after acute stress. This suggests a shift from a combination of glycogenolysis and gluconeogenesis to solely the latter with acute stress. Glucose levels were also associated with uric acid indicating that birds obtain GLU during stress from gluconeogenesis of mostly fat-derived substrates. This provides a previously elusive functional link between body condition and the stress response, and suggests songbirds are more susceptible to stress challenges during energy-limiting conditions than mammals.

## Introduction

An organism’s response to stress (i.e., a challenge to homeostasis) requires the ability to mobilize energy. In vertebrates, a hallmark of the stress response is the secretion of glucocorticoids (e.g., corticosterone or CORT in birds) from the adrenal glands. During acute stress under normal metabolic conditions or during the early stages of fasting (phase I), glucocorticoids stimulate glycogenolysis in concert with catecholamines, glucagon, and growth hormone (reviewed in [1]) and induce lipolysis (e.g., triglyceride (TRIG) breakdown), which results in the release of free glycerol (FG) and free fatty acids (FFAs) ([2]. Free glycerol serves as a substrate for hepatic gluconeogenesis, thus facilitating hyperglycemia after glycogen depletion [2], although glucocorticoids can also promote glycogen deposition when in an absorptive metabolic state [3,4,5]. Basal (i.e., non-stress-associated) glucocorticoids also induce feeding behavior [6,7,8,9,10], but elevated levels suppress appetite [11]. The hyperphagic action of glucocorticoids often elevates plasma TRIG [12] and can counteract the depletion of fat stores due to lipolysis [1]. The state-dependent actions (basal vs. stress) of glucocorticoids are mediated by two distinct glucocorticoid receptor types [13,14,15,16].

During fasting, glucocorticoids stimulate gluconeogenesis and vertebrates rely increasingly on lipid (phase II fasting) and protein catabolism to gain energy, with the latter becoming prominent as fat stores are used up (phase III fasting) [17,18]. Increased amino acid availability due to protein catabolism promotes hepatic enzyme synthesis and glycogen production [19]. Protein catabolism also elevates plasma uric acid (URIC), the predominant nitrogenous end product of protein metabolism in birds and an important antioxidant [20,21]. Sustained and severe fasting, however, elevates the risk of exhausting GLU reserves. This risk is mitigated by promoting the conversion of FFAs to ketones such as β-hydroxybutyrate (β -OHB), that serve as an alternative energy source, particularly to sustain the activity of the nervous system [22,23,24]. Investigating simultaneous changes in plasma concentrations of multiple metabolites during acute stress enables us to infer the energy sources (GLU, lipid, or protein) that animals preferentially use during short-term challenges to homeostasis.

Birds are exceptional models for understanding metabolic changes during acute stress. Compared to size-matched mammals, birds have higher metabolic rates and elevated blood GLU and FFA levels [17], and the energetic demands of flight is fueled by an increased reliance on lipid oxidation for energy [25]. Thus birds respond to a short-term decrease in energy availability faster and with greater loses of body mass than comparably sized mammals [26]. Plasma metabolite profiles track changes in body mass and fluctuate during periods of fasting and energy mobilization [27,28]. The amount of stored energy reserves (i.e., body condition) may, therefore, predict the substrates used during an acute stress response.

Subtle shifts in complex metabolic pathways are difficult to detect using univariate analyses due to intricate and often poorly understood relationships between variables of interest. This issue can be addressed using path analysis, which tests associations between variables based on a hypothetical framework of cause and effect interactions [29,30]. This approach separates the direct and indirect associations between variables and can inform on the specific pathways involved [31,32]. Based on our current understanding of mammalian metabolic pathways, we used path analysis to model how plasma CORT levels alter energy usage, as measured through changes in plasma metabolite (GLU, TRIG, FG, β -OHB, and URIC) levels in four free-living songbird species under both normal (*hereafter* baseline) conditions and after 30 minutes of restraint stress (*hereafter* acute stress), when CORT has increased in circulation.

We tested the hypothesis that the increased energy demands of songbirds during short-term acute stress drive them to primarily mobilize fat and protein stores, thus providing a functional link between CORT and body condition. We studied free-living individuals of four species: two pairs of closely related species from the family Mimidae that are both insectivorous and frugivorous: the Northern Mockingbird (*Mimus polyglottos*) and the Curve-billed Thrasher (*Toxostoma curvirostre*), and two granivorous species, the Abert’s Towhee (*Melozone aberti*) and the House Sparrow (*Passer domesticus*). Besides having different diets, these species may differ with respect to the degree to which excess energy is stored as subcutaneous adipose tissue: mockingbirds and sparrows tend to have greater furcular fat deposits than either thrashers or towhees (HBF, *personal observations*). Birds with smaller subcutaneous fat deposits may rely primarily on lean muscle mass for energy. However, there is limited understanding of the degree to which furcular fat deposits reflect whole body fat reserves in sedentary species that have a limited capacity for subcutaneous fat storage. In migratory species or those from wintering latitudes, furcular fat scores may predict total reserves except, when fat score is close to zero [33,34]. To consider how fat storage may influence metabolite use during acute stress, we also assessed how path models may differ between species that accumulate fat deposits and those that do not. We hypothesized that birds with relatively large fat deposits rely more on fat-derived energy during acute stress than birds with small fat stores.

Based on this hypothesis several predictions can be made concerning the use of TRIG and URIC during stress, the sources of GLU production, and how body condition may be related to circulating GLU and CORT levels. First, songbirds spend a substantial portion of the morning feeding to recover from their overnight fast [35,36] and during this time they are in an absorptive state when anabolism exceeds catabolism. Thus we predict that under baseline conditions and as time of day progresses, increased feeding elevates plasma TRIG levels, which are deposited as fat stores and in turn increases body condition. By contrast, during acute stress TRIG are catabolized to FFAs which are oxidized to produce ketones (i.e., β-OHB) for energy. Furthermore, these relationships are predicted to be stronger in species with high (sparrows and mockingbirds) than low (thrashers and towhees) amounts of furcular fat. Second, baseline plasma CORT may maintain glycemia *via* glycogen stores through interactions with other hormones, but gluconeogenesis is predominant during an acute stress response. Thus we predict that baseline plasma CORT has a direct effect on GLU levels, but this effect decreases during acute stress because CORT mobilizes FG for gluconeogenesis and this effect is predicted to be larger in species with more fat deposition (sparrows, mockingbirds). A further prediction is that body condition impacts the change in plasma GLU either by increasing CORT secretion (CORT-mediated) or by increasing FG levels (fat-mediated) during acute stress. Finally, previous studies found changes in URIC levels during acute stress [37] but whether this stems from protein catabolism for energy or depletion with oxidative stress, a byproduct of GLU production, is uncertain. We predict that if the former, URIC levels would be directly related to CORT and would be stronger in species with low amounts of fat deposition (thrashers, towhees). The latter would be demonstrated by a direct association between GLU and URIC and this association would increase in strength during acute stress.

The present study uses novel statistic approaches to investigate metabolic processes in free-living animals in a manner consistent with the complex “network” structure of metabolic processes. Understanding the interaction of body condition and acute stress is vital as environmental change alters food availability and the high energetic demands of birds make them susceptible to the energy-depleting effects of environmental stressors.

## Results

### Species differences in furcular fat deposits

As predicted, species differed in their furcular fat scores with towhees and thrashers having lower fat scores than sparrows and mockingbirds (Kruskal-Wallis *H*: = 5.7283, *p* = 0.001; Fig 1). Both mockingbird and thrasher data contained outlying points, however these were retained in the analysis, as there was no basis for exclusion. Furcular fat scores were not correlated to time of day nor sampling date for any species (all *p* ≥ 0.092).

**Figure 1.**
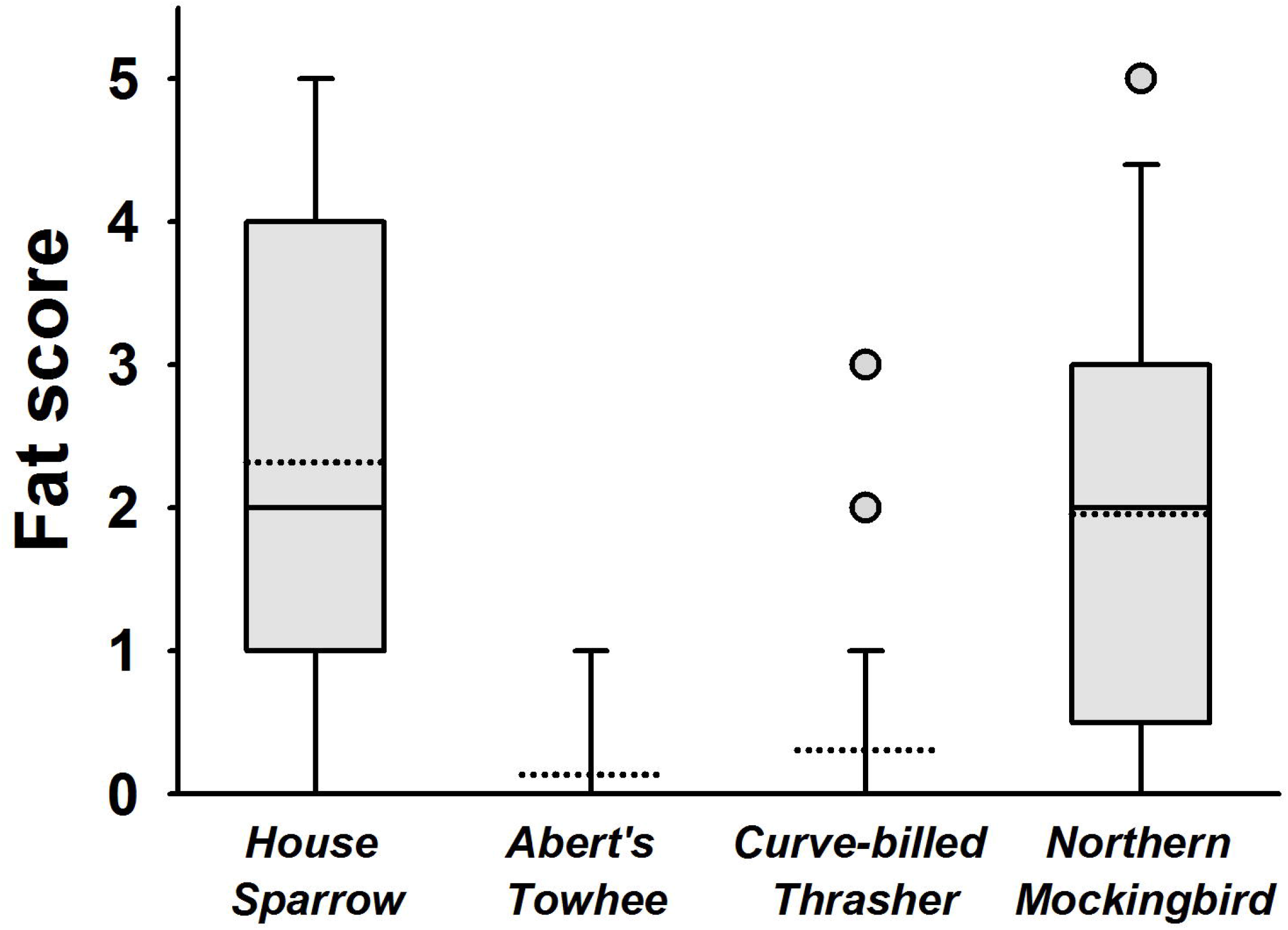
Box plot indicating distribution of furcular fat scores graded on a 5-point scale for four species of songbirds. Boxes represent 25% and 75% quartiles; solid line indicates median; dotted line indicates mean, error bars indicate 5% and 95% confidence intervals and circles indicating statistical outliers (≥ than 2 standard deviations from the mean).

### Baseline and stress-induced plasma hormone and metabolite concentrations

Acute stress increased plasma total and free CORT in all species (Table 1). Acute stress decreased plasma TRIG only in the mockingbird, but plasma FG decreased in three species: towhees, thrashers, and mockingbirds (Table 1). Plasma GLU levels increased with stress in thrashers and mockingbirds, but not in the other two species (Table 1). URIC levels in plasma decreased with stress in all species (Table 1), whereas β-OHB increased with stress only in the towhee (Table 1). Stress did not influence plasma OSMO in any species (Table 1). Sampling date had no statistical effect on plasma CORT or any metabolite in any species (Appendix 1).

**Table 1.** Plasma concentrations of corticosterone (CORT), metabolites, and osmolality (OSMO) for four songbird species at baseline and after 30 minutes of capture and restraint stress. All values indicate mean ± SEM and asterisks indicates significant differences between baseline and stress levels (* *p* ≤ 0.05, and ** Bonferroni corrected *p* ≤ 0.0125).

|  | House Sparrow (N = 40) |  |  | Abert's Towhee (N = 46) |  |  | Curve-billed Thrasher (N = 61) |  |  | Northern Mockingbird (N = 25) |  |  |
| --- | --- | --- | --- | --- | --- | --- | --- | --- | --- | --- | --- | --- |
|  | Baseline | Stress |  | Baseline | Stress |  | Baseline | Stress |  | Baseline | Stress |  |
| **Total CORT (ng/ml)** | 4.16 ± 1.62 | 36.72 ± 6.39 | ** | 12.45 ± 1.35 | 37.22 ± 2.11 | ** | 11.98 ± 2.53 | 57.07 ± 7.97 | ** | 7.63 ± 1.32 | 37.84 ± 8.43 | ** |
| **Free CORT (ng/ml)** | 0.06 ± 0.01 | 0.64 ± 0.10 | ** | 0.19 ± 0.03 | 0.56 ± 0.03 | ** | 0.25 ± 0.07 | 0.93 ± 0.10 | ** | 0.26 ± 0.03 | 0.51 ± 0.11 | * |
| **TRIG (mM)** | 1.99 ± 0.10 | 1.69 ± 0.16 | | 2.01 ± 0.19 | 1.53 ± 0.16 | | 2.89 ± 0.65 | 2.30 ± 0.53 | | 3.02 ± 0.39 | 1.84 ± 0.28 | * |
| **FG (mM)** | 0.76 ± 0.06 | 0.71 ± 0.11 | | 0.86 ± 0.10 | 0.61 ± 0.06 | * | 0.81 ± 0.13 | 0.42 ± 0.06 | ** | 0.97 ± 0.18 | 0.51 ± 0.28 | * |
| **GLU (mM)** | 21.39 ± 3.9 | 20.18 ± 2.62 | | 19.56 ± 0.71 | 18.72 ± 0.87 | | 15.33 ± 0.46 | 18.14 ± 0.68 | * | 17.61 ± 1.21 | 20.08 ± 1.63 | * |
| **URIC (mM)** | 0.37 ± 0.08 | 0.20 ± 0.05 | * | 0.69 ± 0.07 | 0.46 ± 0.05 | * | 1.03 ± 0.15 | 0.52 ± 0.06 | ** | 0.39 ± 0.05 | 0.24 ± 0.05 | * |
| **β-OHB (mM)** | 0.14 ± 0.02 | 0.17 ± 0.03 | | 0.11 ± 0.01 | 0.66 ± 0.02 | ** | 0.12 ± 0.02 | 0.14 ± 0.02 | | 0.07 ± 0.01 | 0.07 ± 0.02 | |
| **OSMO (mOsm/kg)** | 302.69 ± 7.65 | 299.71 ± 4.79 | | 332.50 ± 4.60 | 331.06 ± 4.67 | | 297.62 ± 4.39 | 299.99 ± 4.97 | | 326.70 ± 5.02 | 319.22 ± 5.67 | |

### Model building

Several hypothetical models were tested in each species (Table 2). Major differences between models included the presence or absence of direct relationships between: 1) TRIG and β-OHB; 2) body condition and URIC; 3) time of day and total and free CORT levels; and 4) TRIG and URIC. As these models were tested, non-significant relationships or those with high VIFs (> 10) were removed until the best models were successfully generated for each species and under both baseline and acute stress-induced conditions (Table 2). These best models were then compared using the evaluation statistics described above. Standardized path coefficients are shown for all direct relationships tested across all models and species in Appendix 1. All joint probabilities for the best-supported model family error rates were less than 0.10, suggesting less than a 10% probability of type 1 error.

**Table 2.** Model fit parameters for the three best-supported path analyses models relating time of day, body condition, corticosterone and plasma metabolite levels at baseline and after 30 min of acute stress for four songbird species. Model generally fits to observed data if: 1) χ2 is not-significant, 2) RMSEA is ≤ 0.05; 3) NFI is > 0.9; and 4) the model has a large AIC value.

| Species (No. birds, models tested) | $\chi^2$ | df | *p* | RMSEA | NFI | AIC |
| --- | --- | --- | --- | --- | --- | --- |
| House Sparrow (40, 5) |  |  |  |  |  |  |
| Baseline (*accepted*) | 8.46 | 8 | 0.391 | 0.04 | 0.91 | 74.92 |
| 2nd | 9.73 | 8 | 0.275 | 0.04 | 0.80 | 64.01 |
| 3rd | 9.88 | 7 | 0.302 | 0.05 | 0.78 | 51.30 |
| Stress-induced (*accepted*) | 6.11 | 7 | 0.525 | 0.03 | 0.88 | 62.47 |
| 2nd | 7.91 | 6 | 0.472 | 0.04 | 0.85 | 58.38 |
| 3rd | 8.03 | 5 | 0.451 | 0.05 | 0.83 | 60.32 |
| Abert's Towhee (46, 4) |  |  |  |  |  |  |
| Baseline (*accepted*) | 7.74 | 7 | 0.531 | 0.04 | 0.89 | 68.37 |
| 2nd | 7.32 | 7 | 0.579 | 0.05 | 0.89 | 68.14 |
| 3rd | 8.50 | 7 | 0.310 | 0.06 | 0.83 | 63.96 |
| Stress-induced (*accepted*) | 11.02 | 5 | 0.107 | 0.05 | 0.85 | 80.28 |
| 2nd | 12.75 | 5 | 0.082 | 0.07 | 0.84 | 78.93 |
| 3rd | 10.65 | 6 | 0.134 | 0.06 | 0.92 | 69.23 |
| Curve-billed Thrasher (61, 7) |  |  |  |  |  |  |
| Baseline (*accepted*) | 11.27 | 8 | 0.406 | 0.05 | 0.95 | 88.17 |
| 2nd | 11.01 | 8 | 0.316 | 0.04 | 0.94 | 85.06 |
| 3rd | 8.69 | 7 | 0.260 | 0.06 | 0.95 | 81.35 |
| Stress-induced (*accepted*) | 11.22 | 7 | 0.116 | 0.06 | 0.87 | 87.53 |
| 2nd | 11.19 | 7 | 0.116 | 0.06 | 0.87 | 85.29 |
| 3rd | 13.42 | 7 | 0.089 | 0.07 | 0.82 | 84.15 |
| Northern Mockingbird (25, 5) |  |  |  |  |  |  |
| Baseline (*accepted*) | 12.19 | 7 | 0.247 | 0.06 | 0.97 | 82.13 |
| 2nd | 10.04 | 6 | 0.144 | 0.05 | 0.95 | 86.19 |
| 3rd | 9.55 | 6 | 0.101 | 0.04 | 0.90 | 83.16 |
| Stress-induced (*accepted*) | 10.41 | 7 | 0.238 | 0.06 | 0.98 | 83.11 |
| 2nd | 13.06 | 6 | 0.064 | 0.07 | 0.88 | 62.51 |
| 3rd | 15.32 | 5 | 0.028 | 0.08 | 0.73 | 58.39 |

### Model of baseline interactions

As time of day progressed, baseline TRIG concentrations increase in towhees, sparrows, and thrashers, but not in mockingbirds (Fig 2). Plasma TRIG are positively associated with body condition in all four species and with β-OHB in the thrasher, mockingbird, and sparrow (Fig 2). Body condition is also associated with plasma total, but not free CORT in all four species (Fig 2; Appendix 1). Plasma total CORT is directly associated with FG in three species, and directly with GLU in thrashers and mockingbirds (Fig 2). Similarly free CORT is directly associated with both FG and GLU (Fig 2). The associations are stronger for free than total CORT, with respect to both FG and GLU (Fig 2). URIC levels are negatively associated with plasma GLU in every species (Fig 2).

**Figure 2.**
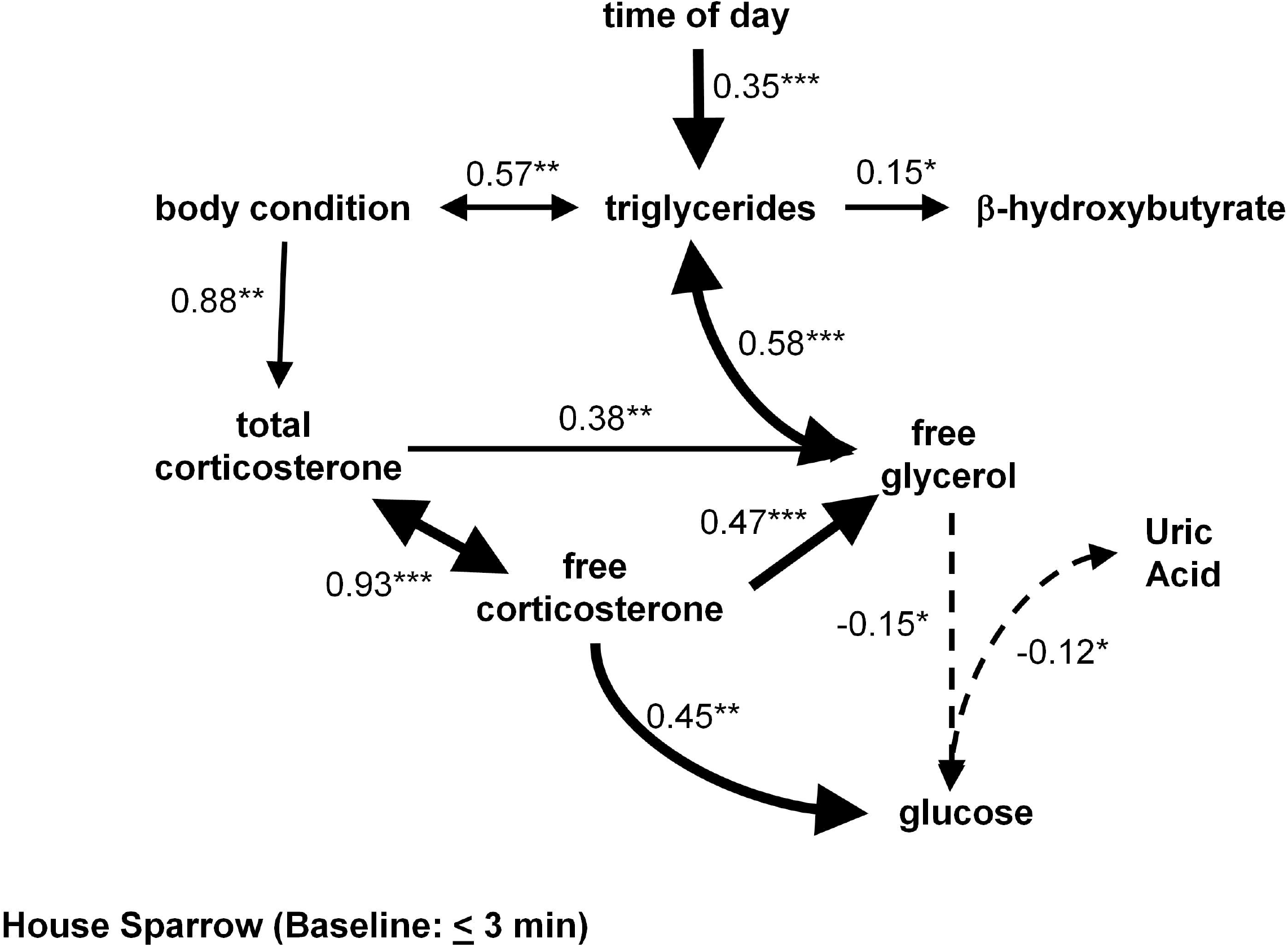
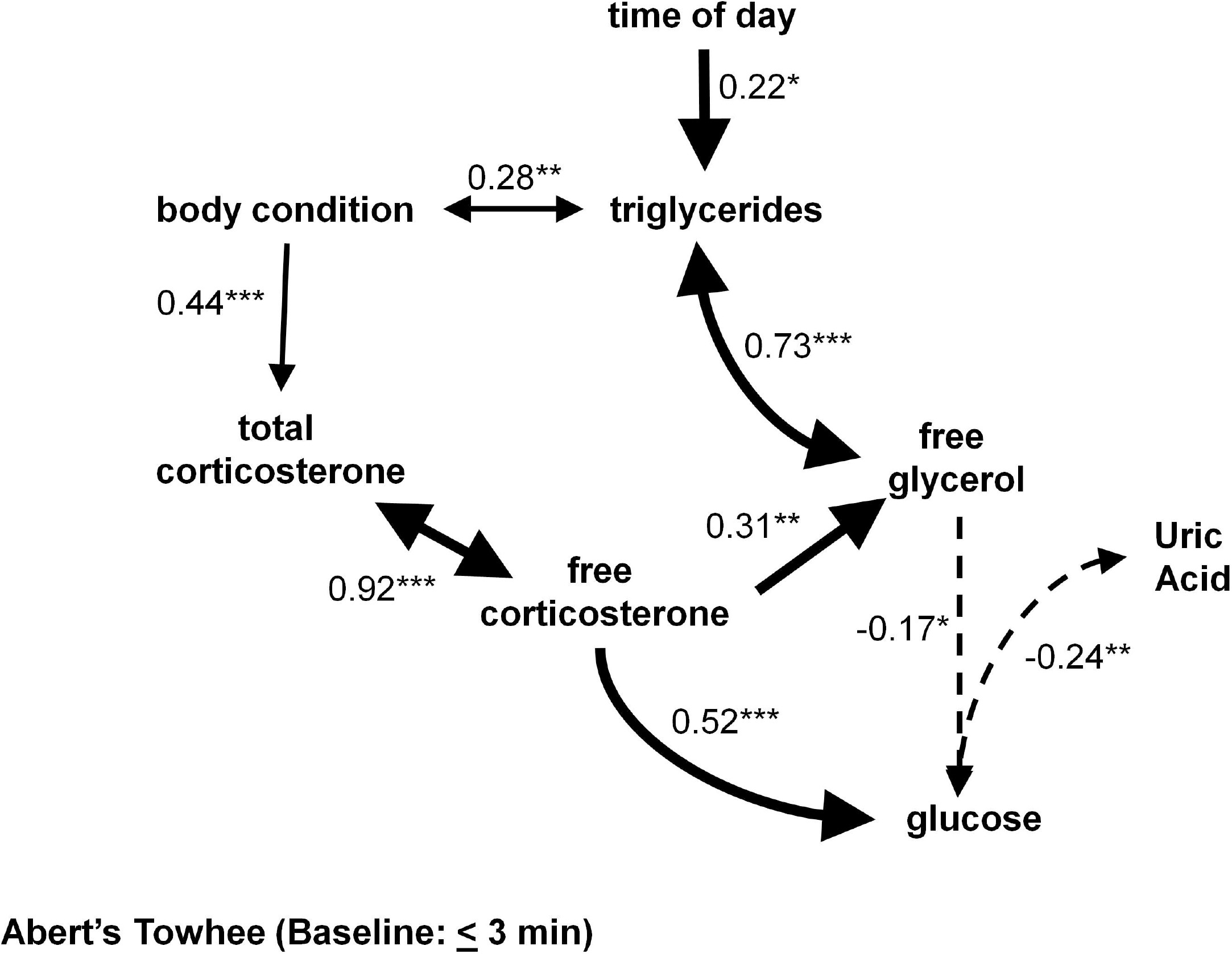
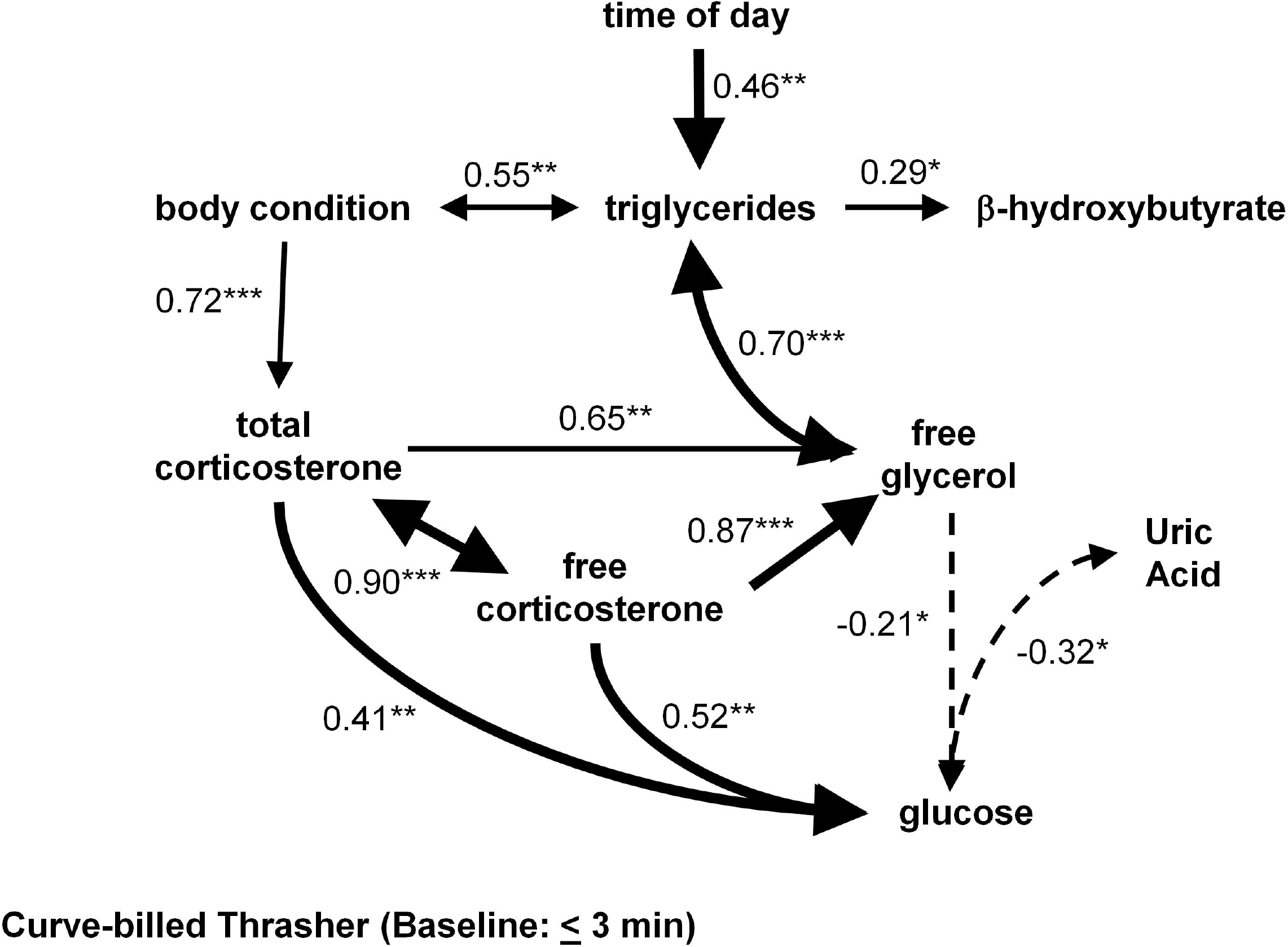
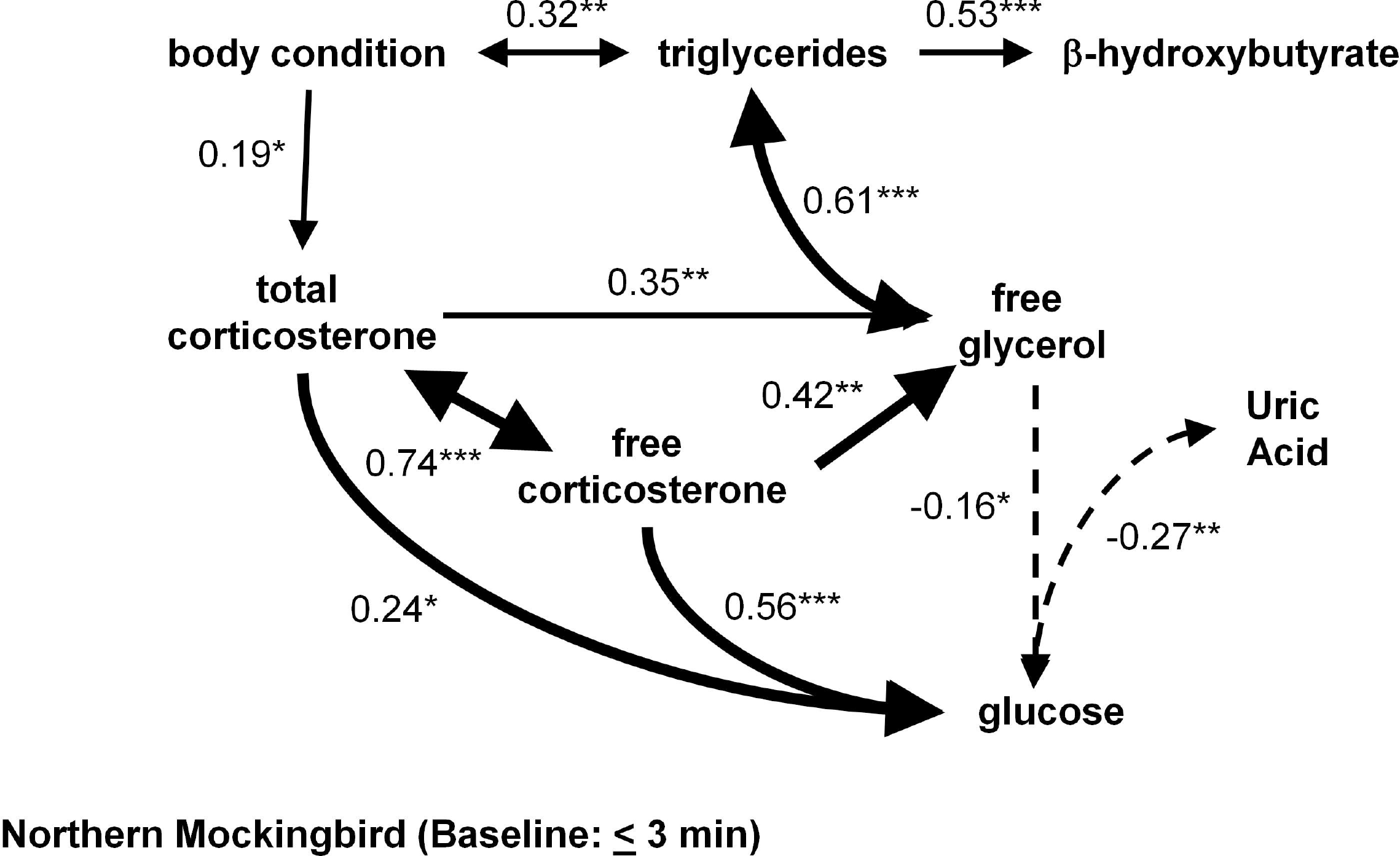
Path model of relationships between plasma metabolites, body condition, and corticosterone during baseline (“unstressed”) conditions for four songbird species: House Sparrows (*Passer domesticus*); Abert’s Towhees (*Melozone aberti*); Curve-billed Thrashers (*Toxostoma curvirostre*); and Northern Mockingbirds (*Mimus polyglottos*). Solid and dashed arrows indicate positive and negative relationships between variables, respectively. Double-headed arrows indicate highly-correlated variables (r ≥ 0.75). Numbers next to arrows represent standardized path (β) coefficients and the thickness of an arrow represents the strength of the relationship between variables. Only arrows for significant relationships are shown, with * indicating a difference at *p* ≤ 0.05; ** indicating a difference at *p* ≤ 0.01; and *** indicating a difference at *p* ≤ 0.001.

### Models of stress-induced changes

Stress-induced TRIG concentrations are positively associated with body condition in all species except towhees (Fig 3). TRIG are also positively associated with β-OHB in sparrows, thrashers, and mockingbirds but this relationship is negative for towhees (Fig 3). Body condition is again positively associated with total, but not free CORT in all four species (Fig 3; Appendix 1). Total CORT is associated with FG negatively (towhees and thrashers) or positively (sparrows and mockingbirds; Fig 3). These relationships are also observed between free CORT and FG concentrations, but the strengths of these path coefficients are greater than for total CORT (Fig 3). Only in mockingbirds do we observe a direct association between CORT (total or free) and GLU levels (Fig 3). Interestingly, significant direct negative associations between plasma FG and GLU concentrations are observed only in thrashers and mockingbirds (Fig 3). FG concentration is also positively associated with β-OHB in towhees and thrashers (Fig 3). URIC levels are again negatively associated with plasma GLU in three species (Fig 3).

**Figure 3.**
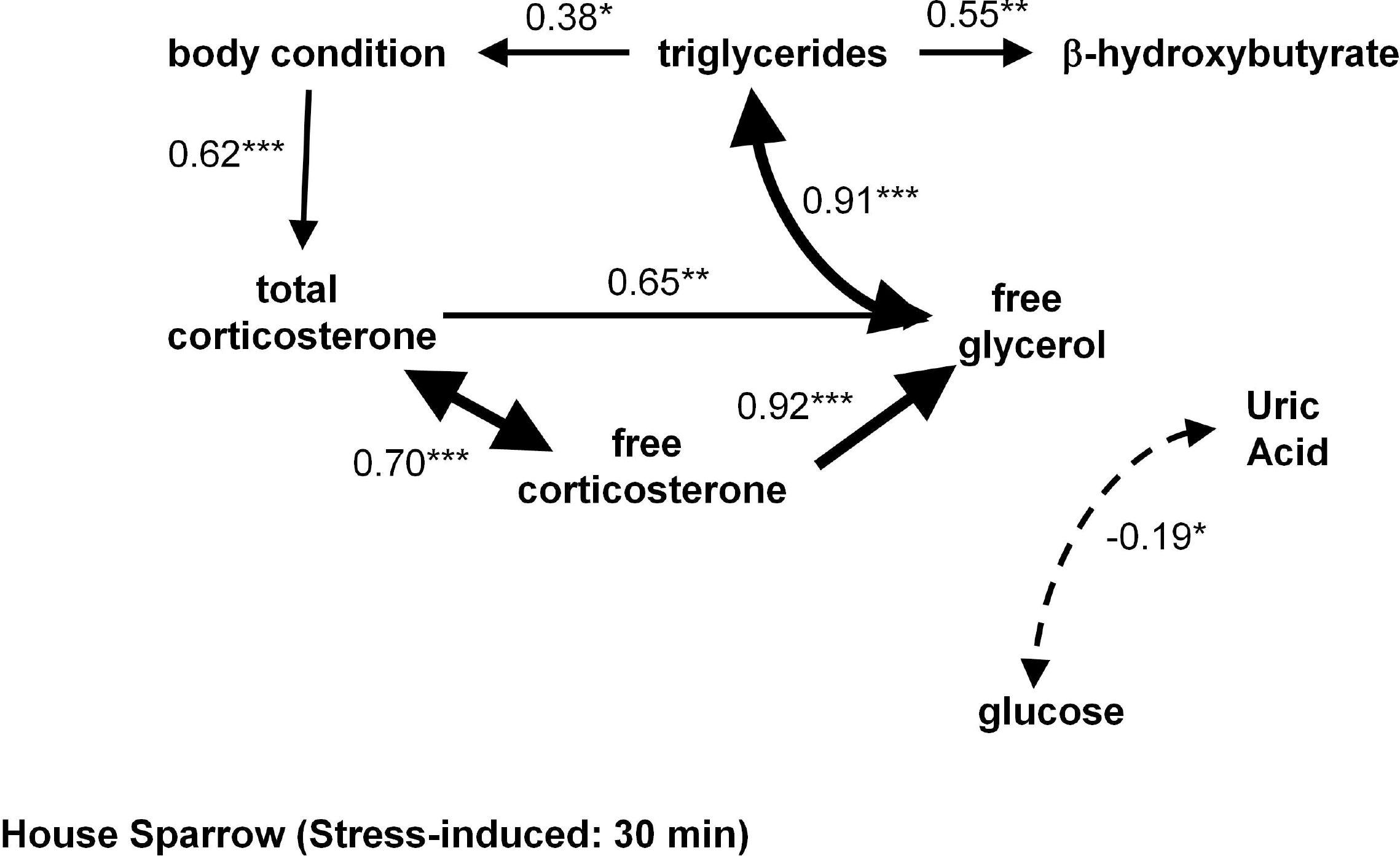
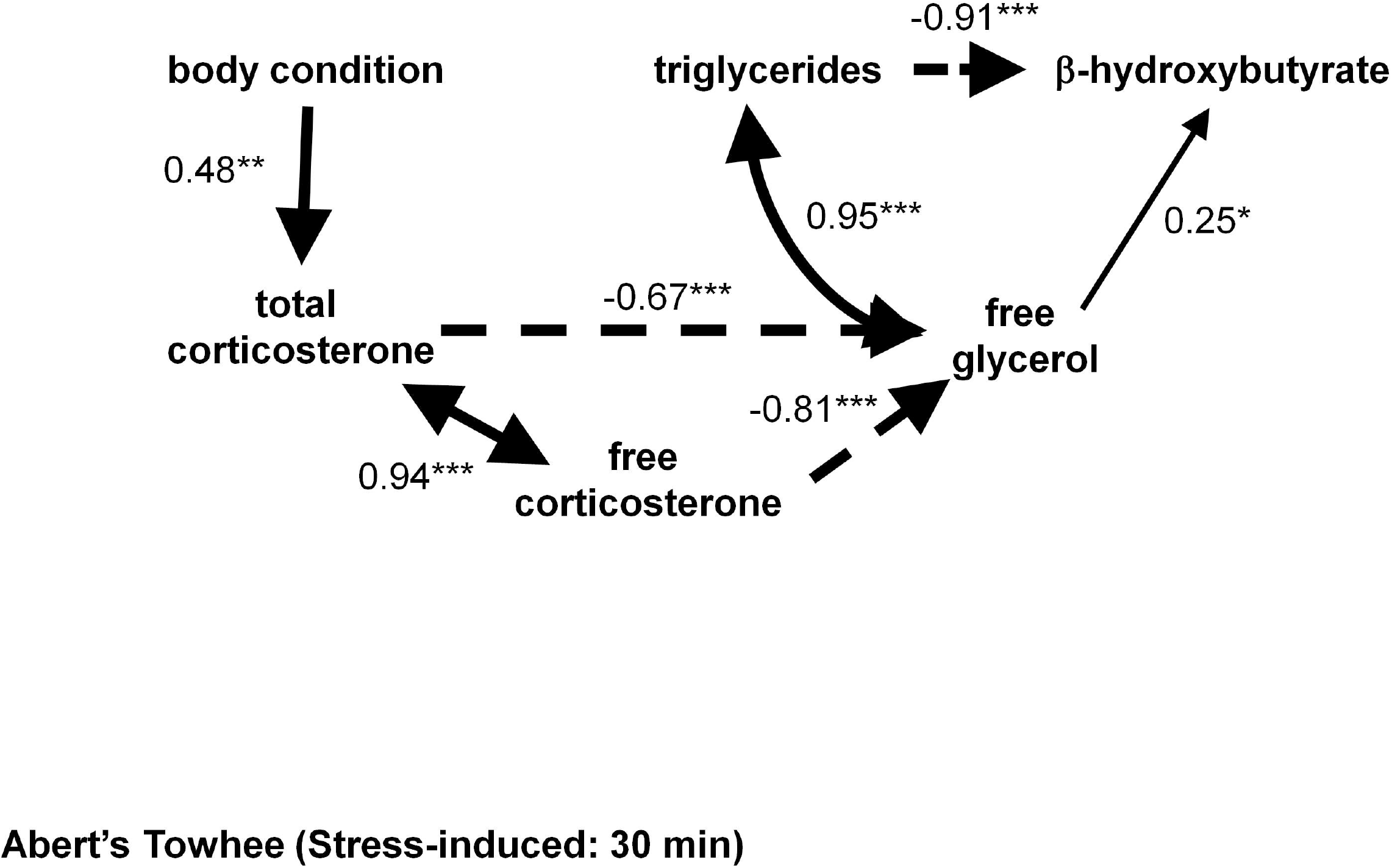
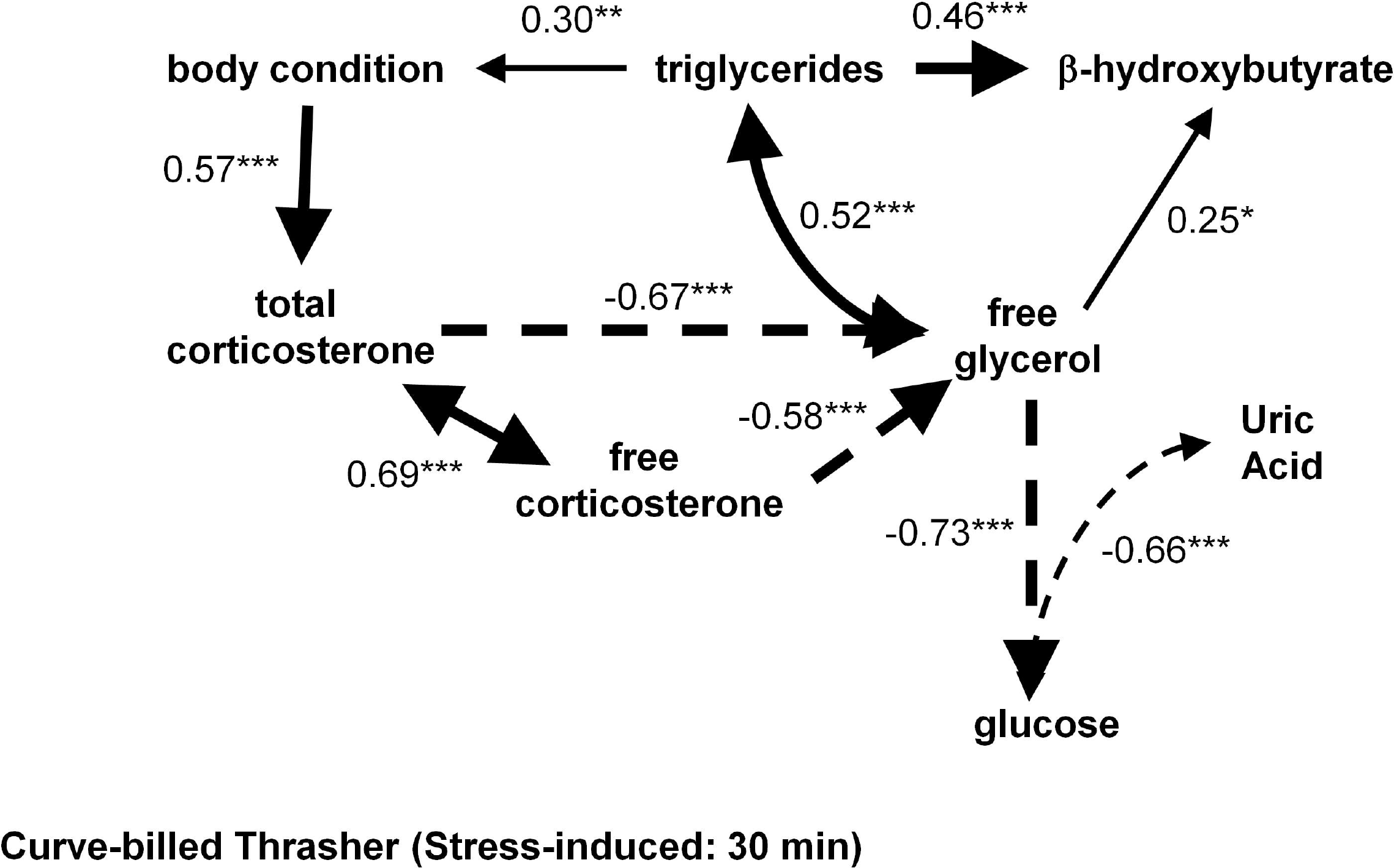
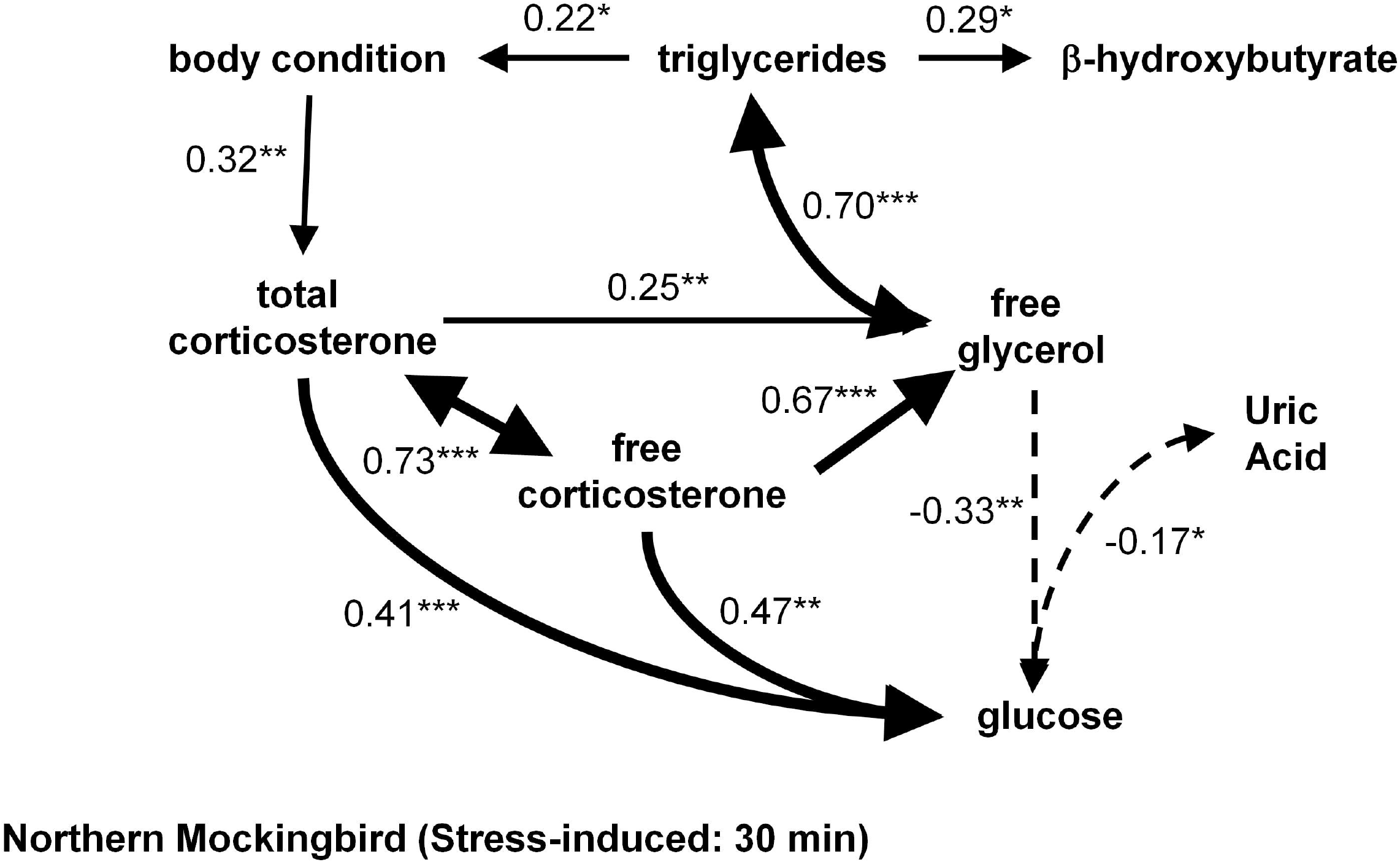
Path model of relationships between plasma metabolites, body condition, and corticosterone after 30 mins of capture and restraint stress for four songbird species: House Sparrows (*Passer domesticus*); Abert’s Towhees (*Melozone aberti*); Curve-billed Thrashers (*Toxostoma curvirostre*); and Northern Mockingbirds (*Mimus polyglottos*). Solid and dashed arrows indicate positive and negative relationships between variables, respectively. Double-headed arrows indicate highly correlated variables (r ≥ 0.75). Numbers next to arrows represent standardized path (β) coefficients and the thickness of the arrow is proportional to the strength of the relationship between variables. Only arrows for significant relationships are shown, with * indicating a difference at *p* ≤ 0.05; ** indicating a difference at *p* ≤ 0.01; and *** indicating a difference at *p* ≤ 0.001.

## Discussion

We used path analysis models to test the hypothesis that during acute stress songbirds derive energy mainly from fat and protein stores. Several models were developed and tested. The most parsimonious models were similar between the four species studied, although some variation in the relative strength and directionality of specific relationships were observed. Thus the metabolic pathways proposed here appear evolutionarily conserved among passerines and they suggest that songbirds increasingly rely on fat, but not protein, for energy during the stress response. Several observations support this conclusion: 1) direct associations between body condition and CORT, 2) a direct association between CORT and GLU present mostly at baseline, 3) switching to associations between CORT and FG during acute stress, 4) a lack of direct associations between CORT or body condition with URIC, and 5) enhanced associations between TRIG and β-OHB with stress in two of four species. Overall this study provides strong precedence for a functional link between body condition and the ability to mount a stress response. Furthermore it provides promise for the use of path analyses to identify general patterns of energy use during complex metabolic events.

As time of day progressed, baseline plasma TRIG levels increased in three species, likely resulting from its accumulation with daily foraging [28,53,54]. Higher plasma TRIG levels indicate fat deposition, which presumably translates to a positive association between TRIG and body condition, as was observed in all species. Despite the increased associations between time of day, condition and TRIG, there was no apparent change in furcular fat score with date or time of day. Furcular fat depots are thought to assist as energy storage sites during migratory flights and for overnight survival, particularly in colder climates [60,61,62]. One interpretation is that furcular fat depots may represent storage sites for fat accumulated at an excess (i.e., well above the required amounts to maintain homeostasis), and these reserves do not change with short-term and small-scale adjustments in body condition. Thus using TRIG levels, amounts of muscle, or a morphometry-based condition index may be better predictors of small-scale metabolic changes in body condition. Interestingly, time of day did not influence baseline TRIG levels in the mockingbird, despite this species having a large furcular fat score. Furthermore, the time of day was not related to TRIG levels during acute stress for any species, suggesting that short-term changes in circulating TRIG levels induced by the stress response are derived from stored sources and not from recently consumed food. Catabolism of these TRIG can provide FG and FFAs as substrates for oxidation to gain energy [25].

All species (except towhee) showed a positive association between baseline TRIG and β-OHB levels, and furthermore all species showed this association during acute stress. Interestingly, in the towhee during stress this association was negative, unlike in other species. Ketones generated by FFA oxidation provide alternative energy substrates for muscle and nervous tissue [24,63]. Higher circulating TRIG may increase background FFA oxidation and thus β-OHB levels, and together may indicate a reliance on ketones as an energy source to supply these tissues [64,65,66]. In two species (thrashers and sparrows), stress increased the strength of the association between TRIG and β-OHB, suggesting greater FFA oxidation as a pathway for supplying energy rapidly. However in towhees, stress generated a negative association between TRIG and β-OHB, whereas in mockingbirds the strength of the association declined with stress. These observations suggest that FFA may not be oxidized during the acute stress response in these species. Alternatively, FFAs in towhees and mockingbirds may increase with stress but are used up prior to the 30 minute sampling point, resulting in no net increase in the concentration of these metabolites being detected. Little is understood concerning the metabolism of ketones in non-mammalian species and birds may serve as fruitful models for future studies on this subject.

We attempted to differentiate between the short-term metabolic effects of plasma total and free CORT. In all species body condition was positively associated with total but not free CORT at baseline and during stress. However, free CORT was in all cases more strongly related (i.e., larger path coefficients) to FG than total CORT. In addition, in all species stress resulted in a stronger association between these variables, although the direction of the relationship differed across species. Free CORT refers to the portion of the circulating steroid capable of diffusing into cells and binding to intracellular receptors [47,67]. There remains considerable debate concerning the relative biological importance of plasma free vs. total CORT [68]. Most studies attempting to differentiate the effects of free and total CORT investigate chronic (days) rather than acute (minutes) roles of CORT [68,69,70,71]. It has been suggested that plasma CBG decreases during acute stress and this decrease could elevate plasma free CORT levels [67]. In previous studies we did not, however, observe such a decrease in any species studied here [39]. Although total and free CORT are correlated, the present observations support the free hormone hypothesis [48] and suggest that short-term changes in free CORT influence the catabolism of TRIG into FG components for gluconeogenesis. Thus, free CORT may reflect the role that this hormone plays in mediating acute metabolic changes during stress more reliably than does total CORT.

In all species at baseline free CORT levels were positively related to FG, but not necessarily to TRIG. In mammals, baseline glucocorticoids mediate fat synthesis and glycogen formation by interacting with insulin in hepatic and adipose tissues [9,72,73]. The consistent relationship between free CORT and FG in this study may reflect a “background” level of CORT-dependent TRIG catabolism that may be necessary to fuel gluconeogenesis in order to maintain baseline blood GLU levels. During acute stress, plasma FG concentrations declined in two species (towhees and thrashers), an observation consistent with previous bird research [28,53]. Other studies, however, reported either an increase [32] or no change [12], and both negative (thrashers and towhees) and positive (mockingbirds and sparrows) associations between stress-induced CORT and FG levels were observed in this study. As the most closely related species (thrashers and mockingbirds) showed opposite relationships between CORT and FG, these results are unlikely explained by phylogeny. In contrast to Sonoran Desert-dwelling towhees and thrashers, wide-ranging and cosmopolitan mockingbirds and sparrows had larger furcular fat stores. In these species, these fat deposits may sustain FG levels during stress resulting in net increases in circulating levels. By contrast, birds with smaller fat reserves (thrashers and towhees) may show a net depletion of plasma FG during stress. This depletion may be exacerbated by the rapid uptake of glycerol by the liver and kidneys, the major avian gluconeogenic sites [74]. However, the degree to which peripheral (e.g., furcular) fat stores sources vs. hepatic TRIG stores are utilized to fuel gluconeogenesis remains unstudied in birds.

Baseline plasma CORT was directly related to GLU levels (i.e., bypassing FG) in all species. At baseline, CORT assists in the regulation of GLU levels by encouraging glycogenolysis [9,72,73]. In response to acute stress, mockingbirds, but not the other species, retained a direct relationship between CORT and GLU. The uncoupling of GLU and CORT supports the hypothesis that stress-induced increases in GLU result from gluconeogenesis with FG as a primary energetic substrate.

However, during stress only two species (mockingbirds and thrashers) demonstrated negative associations between FG and GLU with declines in the former, associated with increases in the latter. In contrast to these two related species, sparrows and towhees showed no significant associations between FG and GLU. GLU did not increase with stress in these species, unlike in thrashers and mockingbirds. One explanation for a lack of hyperglycemia with stress may be the 30 min time frame not being long enough for gluconeogenesis to increase GLU levels. Supporting this contention, plasma GLU in towhees is elevated after one hour, but not after only 30 min of acute stress (Davies and Deviche, *unpublished data*). Alternatively, gluconeogenesis and GLU utilization may occur concurrently, resulting in no net change in circulating GLU levels.

In fasted mammals plasma GLU increases during stress as a result of numerous concurrent effects of glucocorticoids (reviewed in [75]). Data on this subject for birds are less consistent than for mammals. For example, continuous infusion and bolus injections to induce physiological blood CORT levels did not elevate plasma GLU during a 5 hour interval in the turkey (*Meleagris gallapavo*) and produced sporadic increases in plasma GLU in the chicken (*Gallus domesticus*) but only after several hours [76]. In European Starlings (*Sturnus vulgaris*), GLU also did not change with repeated acute stress despite fluctuations in plasma CORT [77], but in another study exogenous CORT induced hyperglycemia during the day but not at night [78]. By contrast, an earlier study on this species demonstrated a hyperglycemic effect of acute stress during the night (when GLU is low) but not during the day (when GLU is high) [79]. Recent food intake may also influence plasma GLU levels. In the present study, birds captured in the morning were likely foraging in an effort to replenish energy reserves used up throughout the night [28]. Birds captured later in the day (i.e., having had more time available to forage and being in an absorptive state) may have higher GLU levels than those captured earlier, but this was not observed. Time of day was associated with increased TRIG (i.e., likely indicating feeding) and thus plasma GLU levels are unlikely to reflect recent feeding activity.

Besides fat stores, birds can also use muscle proteins as substrates for gluconeogenesis [80] as well as for powering flight [81]. We measured changes in levels of URIC, which is derived from the degradation of amino and nucleic acids and can reflect protein (esp. purine) catabolism [82] at baseline and after acute stress. In free-living migrating and captive birds, plasma URIC levels increase when body mass declines [12,83]. However, we did not observe a direct relationship between body condition and URIC in this study for any species. Furthermore, URIC decreased with stress in all species and was negatively associated only with GLU in all models, with the exception of stressed towhees. Decreased URIC levels during acute stress may result from excretion of this metabolite, which is the primary nitrogenous waste product in birds. However, in this study most birds did not appear to have excreted during the 30 min restraint period (*personal observation*), although feces may be retained in the cloaca and not visible to the researcher. Thus circulating URIC levels likely do not reflect mobilization of proteins during stress.

URIC also acts as a potent antioxidant to quench free radicals generated during gluconeogenesis [84,85,86,87] and previous studies have reported on its antioxidant properties in birds [21,37,88]. Birds have higher circulating URIC concentrations than similar-sized mammals [17,84], which may explain their decreased susceptibility to oxidative stress despite having higher GLU levels [89,90]. In a study of 57 bird species, the majority demonstrated a decline in plasma URIC after one hour of acute stress [88]. A rapid stress-associated increase in plasma GLU due to gluconeogenesis may induce oxidative stress which may deplete URIC to prevent oxidative damage to tissue. This hypothesis is speculative but the antioxidant properties of URIC are not well understood and thus deserve further experimental study.

Due largely to their increased energy demands, songbirds respond metabolically to acute stress to a greater degree than mammals. Their physiological responses to energy limitation and stress are more robust than the modest responses of mammals and avian species can, therefore, serve as excellent models to investigate the metabolic effects of stress. In addition, the ability of birds to rapidly mobilize fat resources to maintain glucose levels during stress makes them particularly susceptible to stress when their body condition is already compromised. The use of modeling to assess changes in metabolites may help assess health condition at the individual and population levels.

## Materials and methods

All procedures were pre-approved by the Arizona State University Institutional Animal Care and Use Committee and were conducted with appropriate permits from the Arizona Game and Fish Department, Bureau of Land Management, United States Fish and Wildlife Service, United States Geological Survey’s Bird Banding Laboratory, and the Parks and Recreation Departments for the cities of Phoenix, Tempe, Scottsdale, and Laveen. Also all private landowners first granted us permission to access their lands for sampling. No endangered or protected species were used in this study

### Sampling locations

Birds were sampled in and around Phoenix, Arizona (USA) and locations varied between species, but included a mix of urban-suburban (all species), and either farmlands (sparrows) or undeveloped Sonoran Desert habitats (mockingbird, thrasher, and towhee) with all locations no more than 25 km apart. Detailed information on study sites is found in [38,39]. Previous research suggested that urban birds are often in better body condition and have greater CORT responses to stress than conspecifics from desert areas [38,39]. Individuals from all sampling locations were analyzed together to provide data over a wide range of body conditions.

### Field data collection

Birds were caught using mist nets and either passively (sparrows) or with conspecific song playback recordings (other species), and within 5 hours of sunrise. Previous research demonstrated no effect of song playback on CORT levels [40]. Sample sizes were as follows: mockingbirds: *N* = 25; thrashers: *N* = 61; towhees: *N* = 46; and House sparrows: N = 41. Sampling occurred between March and June 2006 and between March and May 2007. These periods coincided with the breeding season of all birds ranging from incubation to chick-rearing. Previous research did not demonstrate changes in plasma CORT within these periods [39]. Nonetheless sampling date was included in model building to test for seasonal changes in metabolite levels. The time taken to capture the bird may impact circulating CORT levels and this can potentially alter the metabolic profile. However previous research in these species did not show any relationship between capture time and plasma CORT concentrations in either thrashers or towhees [40] and this was unquantifiable in sparrows that were passively netted. Furthermore, sample size limits the number of variables that can be tested in a path model at one time, thus time of capture was not included in the model [29,30]. Upon capture a blood sample (~200 μl) was taken within 3 min from the right jugular vein using a heparinized 0.3 ml syringe and a 29.5 gauge needle. Plasma CORT did not increase significantly during the first 3 minutes following capture (*data not shown*), although such an increase was previously observed in House Sparrows [41]. These samples were defined as pre-stress (i.e., baseline) samples. Birds were then held in a cloth bag for 30 min and a second blood sample (~200 μl) was collected in the same fashion. This capture and restraint protocol is commonly used to induce an acute stress response [42,43]. Blood samples were kept on ice until centrifuged to separate formed elements from plasma, which was stored at -80 °C until assayed.

Only adult males in breeding condition were used and age, sex, and breeding status were established using plumage characteristics [44] and unilateral laparotomy after the second blood sample was collected [38,39]. Body mass (+ 0.1 g) and wing chord (+ 1 mm) were measured to generate body condition indices. To assess differences in subcutaneous fat stores between species, we scored furcular fat (scale of 1 to 5) according to [44,45,46]. Each bird received a uniquely numbered aluminum leg band and was released at the capture location.

### Plasma CORT Assays

Most plasma CORT circulates bound to CORT binding globulins (CBG) [47] and it is suggested that only the unbound fraction of hormone (*hereafter* free CORT) binds to intracellular receptors in target cells [48,49]. Thus both total (i.e., bound and unbound) and free (i.e., unbound) CORT was measured. Plasma total CORT was measured using commercial competitive enzyme-linked immunoassays (ELISA; Assay Designs Inc., Ann Arbor, Michigan, USA) as described and validated by [39]. Samples were assayed in duplicate and with baseline and stress-induced samples from the same individual assayed on the same plate. The sensitivity of the assay ranged from 5.8 -18.1 pg/ml depending on the plate and the mean intra- and inter-assay coefficients of variation were 8.6%, and 14.7%, respectively. Radioligand binding assays were used to estimate plasma CBG binding capacity according to [50,51] with minor modifications specified in [39]. Previously described equilibrium dissociation constants for CORT binding to CBG for each species [39] were used to estimate plasma concentrations of free and CBG-bound CORT according to the equation of [52].

### Plasma Metabolite Assays

Plasma FG and TRIG were measured using a sequential color endpoint assay (Sigma-Aldrich, reagents F6428 and T2449) modified for small plasma volumes (5 μl) and described in [28,53]. Plasma GLU and β-OHB were measured using colorimetric enzyme endpoint assays (Cayman Chemical Co. Ann Arbor, Michigan, USA; Cat No. 10009582 and 700190, respectively). Plasma URIC was also measured using a colorimetric assay (Biovision Research, Mountain View, California, USA; Cat No. K608-100). Previous studies have validated these assays for use in several songbird species [2,27,28,54]. Samples were assayed in duplicate and in random order, and all concentrations are expressed in the same unit (mM) to facilitate comparisons. Assay sensitivities and mean intra- and inter-assay coefficients of variation are as follows: FG: 0.05 – 6.3 mM, 6.5% and 13.0%; TRIG: 0.07 – 11.4 mM, 7.2% and 10.1%; GLU: 2.5 – 28.36 mM, 3.3% and 11.9%; β- OHB: 0.01 – 4.8 mM, 5.5% and 14.0%; and URIC: 0.01 – 6.1 mM, 3.7% and 9.8%.

To determine whether stress-induced biochemical changes alter plasma solute concentration, plasma osmolality (OSMO: mOsm/l) was measured using a vapor pressure osmometer (Model 5500XR, Wescor Inc. Logan, Utah, USA) with 10 μl samples assayed in duplicate. The osmometer was calibrated to known concentration standards (290 and 1000 mOSm/kg) before use, and intra-assay sensitivity was 3.4%.

### Path analysis

Pairwise differences between baseline and stress-induced concentrations of metabolites and CORT were assessed using paired Student’s t-tests. Linear regressions of body mass on wing chord provided standardized residuals for use as body condition indices for each species. Species differences in furcular fat scores were assessed using Kruskal-Wallis analysis of variance (ANOVA), and effects of time of day and date were assessed using Pearson correlations. When necessary, continuous data were natural logarithm (ln) transformed to satisfy normality assumptions. Path analysis enables testing of an *a priori* hypothesis of relationships generated on the basis of theory and the results of Pearson’s correlation analysis between variables. The direction and strength of relationships between each variable was quantified using path (β) coefficients calculated using the maximum likelihood method [30]. The effects of factors outside the model, such as measurement errors, were represented by residual error terms (i.e., unexplained variation) in the model. Predictor variables that are highly correlated (i.e., collinear) can inflate the variance of β coefficients and decrease their accuracy from the “true” value [55]. The effects of collinearity on β coefficients is difficult to detect and can be estimated by examining variance inflation factors (VIFs) for each predictor variable. A VIF of less than 10 is generally considered acceptable [56]. Hypothetical models (or representations) were compared to observed data to determine the goodness of fit. Non-significant relationships (P ≥ 0.05) between variables were removed from the path diagrams, except where necessary to maintain connectivity, and until the most parsimonious and best-supported model was attained.

The multiple comparisons associated with path analytical model building increase the likelihood of type 1 errors, but conventional multiple comparison corrections (e.g., Bonferroni) would be far too conservative to enable model development. Instead the joint probability, a multiple comparison significance test specifically designed for path analysis, was calculated [57]. This method provides the probability of a type 1 error being committed using all significant tests in a model (i.e., family error rate) or in specific number of comparisons using critical alpha values generated by the path model.

Evaluation of alternative models involved several approaches. The hypothetical models were tested using a χ^2^ goodness of fit test statistic, in which a smaller value indicates better consistency with observed data. The root mean square error of approximation (RMSEA), which estimates the amount by which estimated values differ from actual values, was also used for model comparison. A RMSEA ≤ 0.05 is usually considered to indicate a ‘close fit’ to the observed data [58]. To eliminate spurious relationships, the hypothetical model was tested against a saturated model in which all variables are directly connected to each other, and against an independent model (equivalent to a traditional multiple regression) in which no connection exists between variables. The Normed Fit Index (NFI) compares the hypothetical model to both the saturated and independent models. Larger NFI values are preferred and values > 0.9 are generally considered adequate [59]. The Akaike information criterion (AIC) also distinguishes between models derived from the maximum likelihood with the most parsimonious model being associated with the smallest AIC value. Together, these tests provide a rigorous assessment and comparison of different models. For brevity, only model-building results for the top three parsimonious models are presented for each species. Statistical analyses were performed using SPSS Version 13.0 (2004) with the AMOS 7.0 extension for path analysis.

## Acknowledgements

We appreciate many generous landowners, businesses, and agency personnel for providing access to sampling locations. We especially thank Pierre Deviche, Karen Sweazea, Miles Orchinik, Dale DeNardo, Kevin McGraw, Ananias Escalante, Laura Hurley, and Scott Davies, for financial and logistical support and comments on earlier versions of this preprint. This study was supported by National Science Foundation award DEB-0423704 to the Central Arizona–Phoenix Long-Term Ecological Research as well as additional funds to HBF from the Society for Integrative and Comparative Biologists, Arizona State University (ASU) Graduate and Professional Student Association, ASU School of Life Sciences, ASU chapter of the Sigma Xi Scientific Society, and the Natural Sciences and Engineering Research Council.

## Appendix 1.

Relationships tested using path analysis between time of day, date, and plasma levels of metabolites and corticosterone (CORT) in four songbird species at baseline and after 30 mins of capture and restraint stress. Shown are standardized path coefficients (β) and P-values. Significant relationships (p ≤ 0.05) are shown in **bold.**

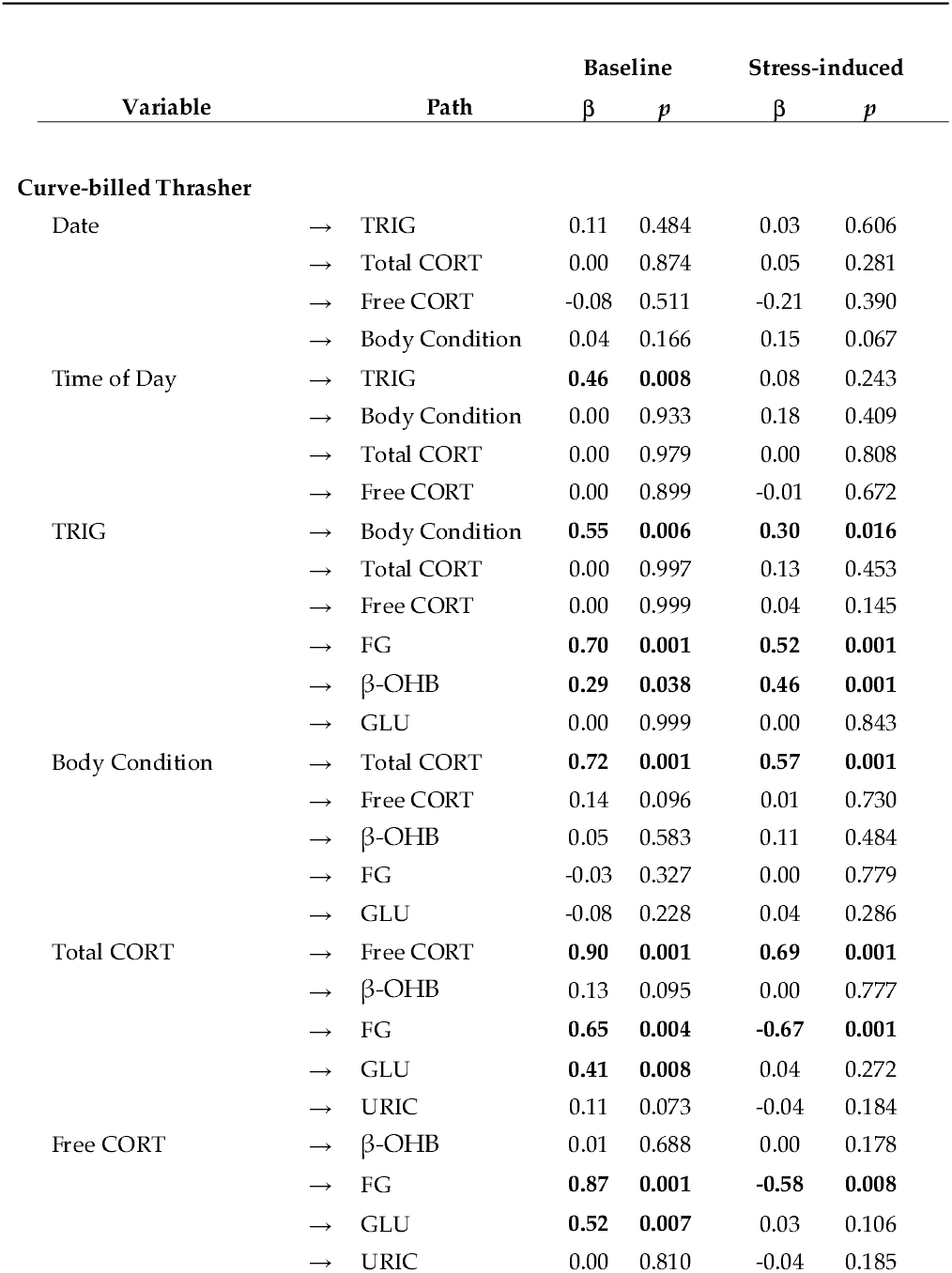
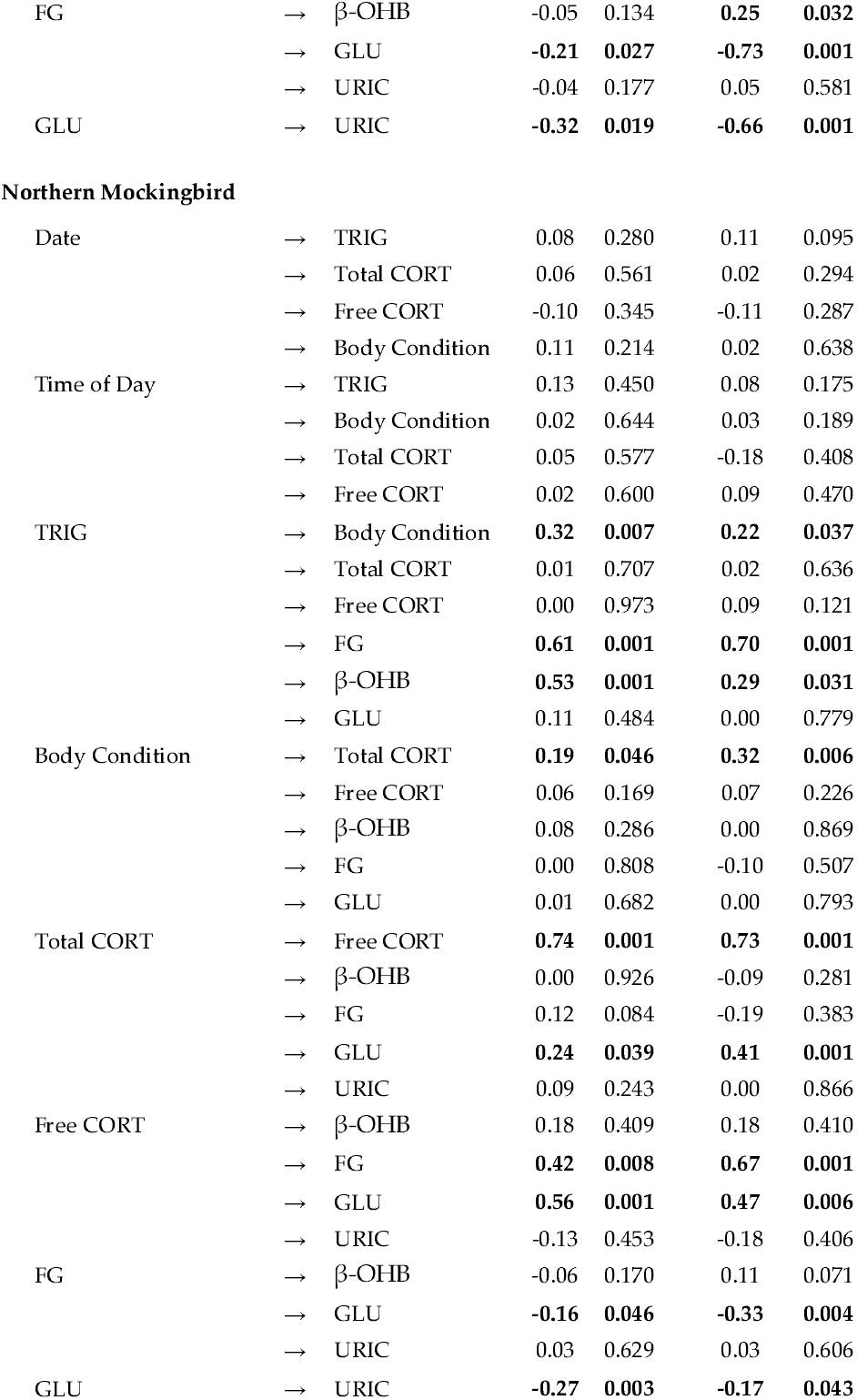
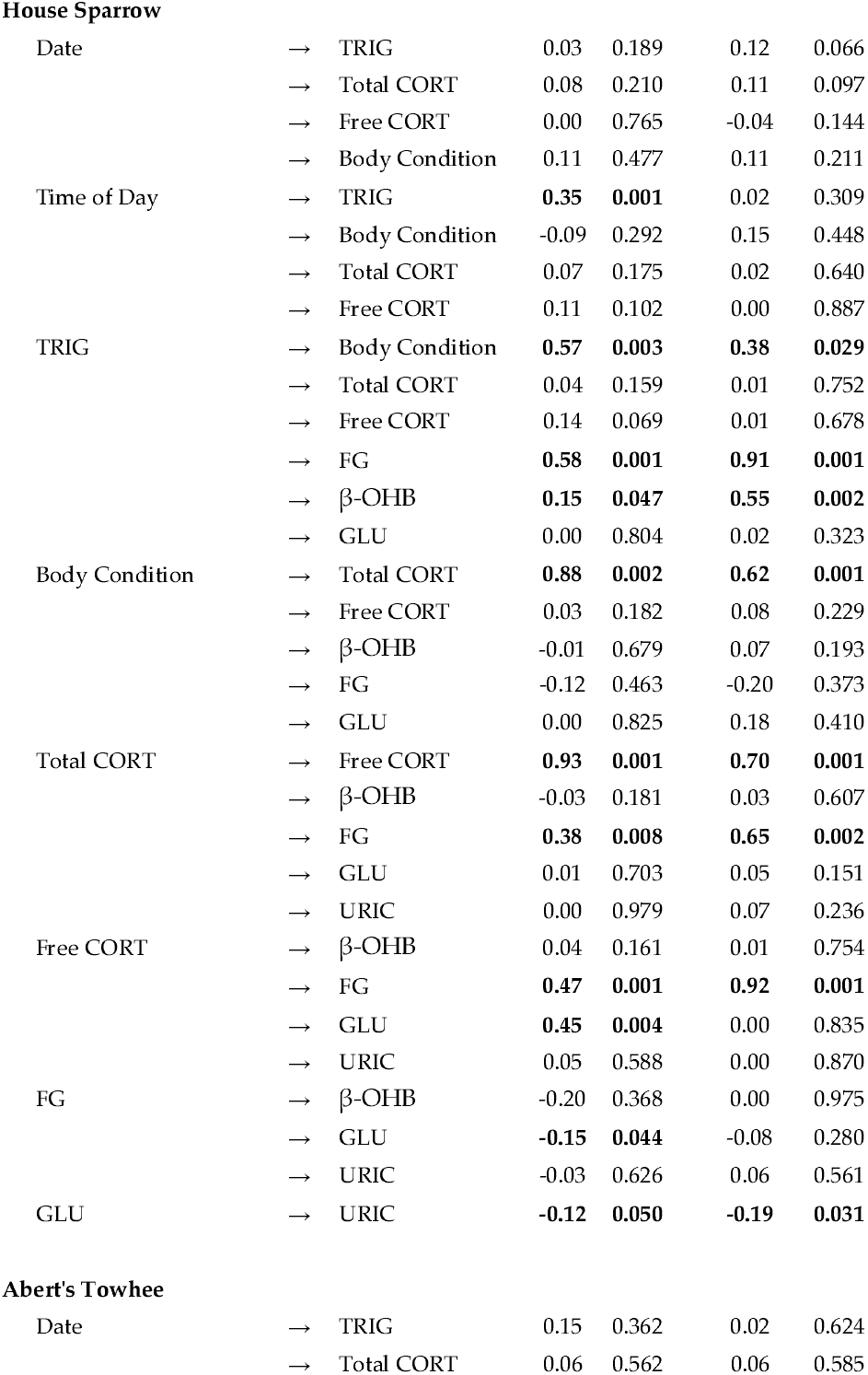
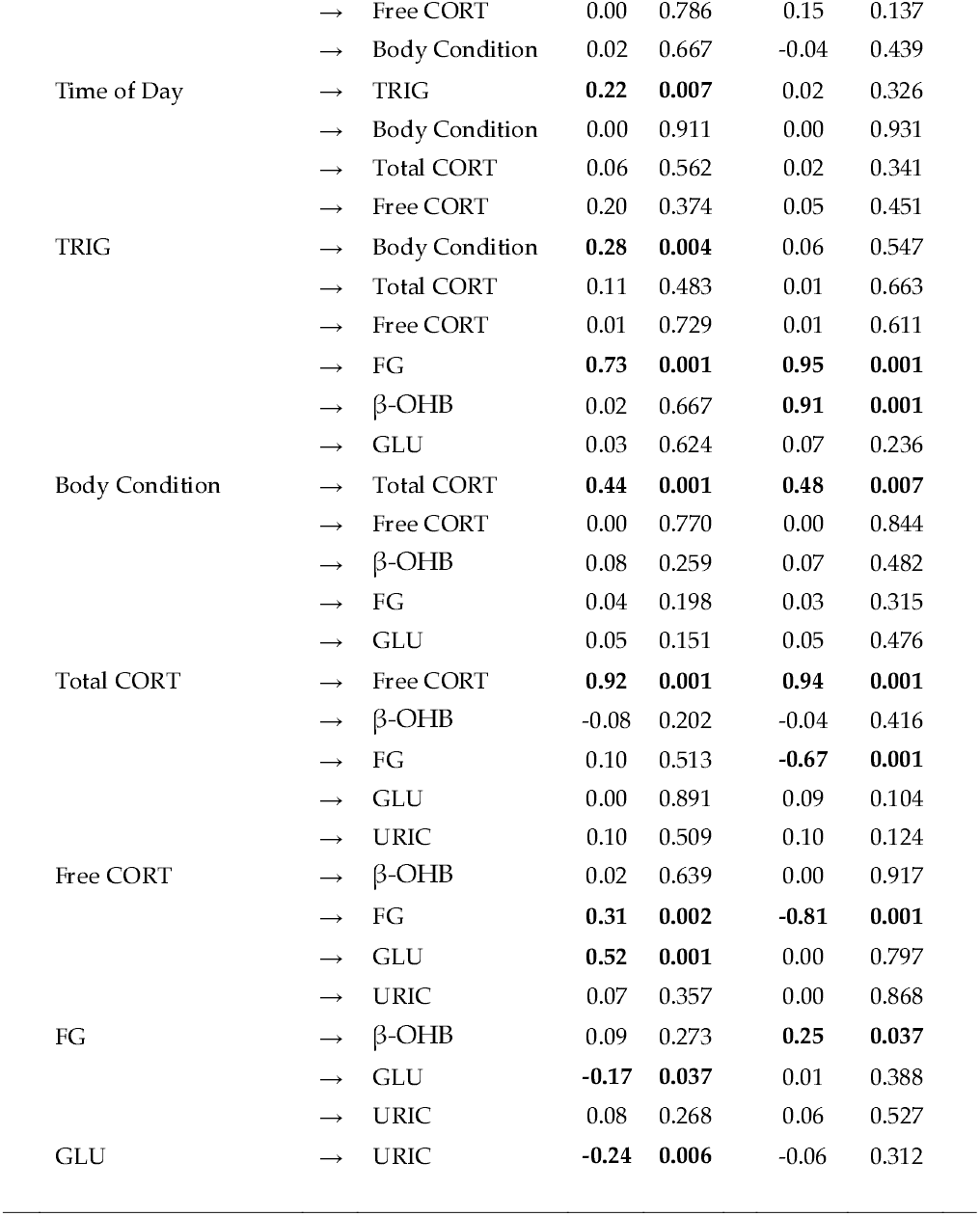

## References

1. Sapolsky RM, Romero LM, Munck AU (2000) How do glucocorticoids influence stress responses? Integrating permissive, suppressive, stimulatory, and preparative actions. Endocrine Reviews 21: 55–89.

2. Jenni-Eiermann S, Jenni L, Piersma T (2002) Plasma metabolites reflect seasonally changing metabolic processes in a long-distance migrant shorebird (Calidris canutus). Zoology 105: 239–246.

3. Bollen M, Keppens S, Stalmans W (1998) Specific features of glycogen metabolism in the liver. Biochemical Journal 336:19–31.

4. Cardell RR (1974) Morphological actions of glucocorticoids on glycogen deposition in liver cells. Anatomical Record 178: 320–321.

5. Zheng XF, Liu L, Zhou J, Miao MY, Zhou JR, et al. (2009) Biphasic effects of dexamethasone on glycogen metabolism in primary cultured rat hepatocytes. Journal of Endocrinological Investigation 32: 756–758.

6. Kitaysky AS, Wingfield JC, Piatt JF (2001) Corticosterone facilitates begging and affects resource allocation in the black-legged kittiwake. Behavioral Ecology 12: 619–625.

7. Pravosudov VV (2003) Long-term moderate elevation of corticosterone facilitates avian food-caching behaviour and enhances spatial memory. Proceedings of the Royal Society of London Series B-Biological Sciences 270: 2599–2604.

8. Bates HE, Kiraly MA, Yue JTY, Montes DG, Elliott ME, et al. (2007) Recurrent intermittent restraint delays fed and fasting hyperglycemia and improves glucose return to baseline levels during glucose tolerance tests in the Zucker diabetic fatty rat - role of food intake and corticosterone. Metabolism-Clinical and Experimental 56: 1065–1075.

9. Warne JP, Akana SF, Ginsberg AB, Horneman HF, Pecoraro NC, et al. (2009) Disengaging insulin from corticosterone: roles of each on energy intake and disposition. American Journal of Physiology-Regulatory Integrative and Comparative Physiology 296: R1366–R1375.

10. Foster MT, Warne JP, Ginsberg AB, Horneman HF, Pecoraro NC, et al. (2009) Palatable Foods, Stress, and Energy Stores Sculpt Corticotropin-Releasing Factor, Adrenocorticotropin, and Corticosterone Concentrations after Restraint. Endocrinology 150: 2325–2333.

11. Liu XY, Shi JH, Du WH, Fan YP, Hu XL, et al. (2011) Glucocorticoids decrease body weight and food intake and inhibit appetite regulatory peptide expression in the hypothalamus of rats. Experimental and Therapeutic Medicine 2: 977–984.

12. Seaman DA, Guglielmo CG, Williams TD (2005) Effects of physiological state, mass change and diet on plasma metabolite profiles in the western sandpiper Calidris mauri. Journal of Experimental Biology 208: 761–769.

13. Dekloet ER, Ratka A, Reul J, Sutanto W, Vaneekelen JAM (1987) Corticosteroid receptor types in brain - regulation and putative function. Annals of the New York Academy of Sciences 512: 351–361.

14. Dekloet ER, Reul J, Deronde FSW, Bloemers M, Ratka A (1986) Function and plasticity of brain corticosteroid receptor systems - action of neuropeptides. Journal of Steroid Biochemistry and Molecular Biology 25: 723–731.

15. Landys MM, Ramenofsky M, Wingfield JC (2006) Actions of glucocorticoids at a seasonal baseline as compared to stress-related levels in the regulation of periodic life processes. General and comparative endocrinology 148: 132–149.

16. Reul J, Dekloet ER (1985) Two receptor systems for corticosterone in rat brain - microdistribution and differential occupation. Endocrinology 117: 2505–2511.

17. Braun EJ, Sweazea KL (2008) Glucose regulation in birds. Comparative Biochemistry and Physiology B-Biochemistry & Molecular Biology 151: 1–9.

18. Dong H, Lin H, Jiao HC, Song ZG, Zhao JP, et al. (2007) Altered development and protein metabolism in skeletal muscles of broiler chickens (Gallus gallus domesticus) by corticosterone. Comparative Biochemistry and Physiology a-Molecular & Integrative Physiology 147: 189–195.

19. Coderre L, Srivastava AK, Chiasson JL (1991) Role of glucocorticoid in the regulation of glycogen-metabolism in skeletal-muscle. American Journal of Physiology 260: E927–E932.

20. Fery F, Plat L, Melot C, Balasse EO (1996) Role of fat-derived substrates in the regulation of gluconeogenesis during fasting. American Journal of Physiology-Endocrinology and Metabolism 270: E822–E830.

21. Cohen AA, McGraw KJ, Robinson WD (2009) Serum antioxidant levels in wild birds vary in relation to diet, season, life history strategy, and species. Oecologia 161: 673–683.

22. Castellini MA, Costa DP (1990) Relationships between plasma ketones and fasting duration in neonatal elephant seals. American Journal of Physiology 259: R1086–R1089.

23. Jenni-Eiermann S, Jenni L (2001) Postexercise ketosis in night-migrating passerine birds. Physiological and Biochemical Zoology 74: 90–101.

24. Peters A, Schweiger U, Pellerin L, Hubold C, Oltmanns KM, et al. (2004) The selfish brain: competition for energy resources. Neuroscience and Biobehavioral Reviews 28: 143–180.

25. Jenni L, Jenni-Eiermann S (1998) Fuel supply and metabolic constraints in migrating birds. Journal of Avian Biology 29: 521–528.

26. Fokidis HB, Hurley L, Rogowski C, Sweazea K, Deviche P (2011) Effects of Captivity and Body Condition on Plasma Corticosterone, Locomotor Behavior, and Plasma Metabolites in Curve-Billed Thrashers. Physiological and Biochemical Zoology 84.

27. Williams TD, Guglielmo CG, Egeler O, Martyniuk CJ (1999) Plasma lipid metabolites provide information on mass change over several days in captive western sandpipers. Auk 116: 994–1000.

28. Guglielmo CG, O’Hara PD, Williams TD (2002) Extrinsic and intrinsic sources of variation in plasma lipid metabolites of free-living western sandpipers (Calidris mauri). Auk 119: 437–445.

29. Sinervo B, DeNardo DF (1996) Costs of reproduction in the wild: Path analysis of natural selection and experimental tests of causation. Evolution 50: 1299–1313.

30. Sokal RR, Rohlf FJ (1995) Biometry: the principles and practice of statistics in biological research: Freeman.

31. Kern M, Bacon W, Long D, Cowie RJ (2005) Blood metabolite and corticosterone levels in breeding adult Pied Flycatchers. Condor 107: 665–677.

32. Kern MD, Bacon W, Long D, Cowie RJ (2007) Blood metabolite levels in normal and handicapped pied flycatchers rearing broods of different sizes. Comparative Biochemistry and Physiology a-Molecular & Integrative Physiology 147: 70–76.

33. Rogers CM (1991) An evaluation of the method of estimating body fat in birds by quantifying visible subcutaneous fat. Journal of Field Ornithology 62: 349–356.

34. Spengler TJ, Leberg PL, Barrow WC (1995) Comparison of condition indexes in migratory passerines at a stopover site in coastal Louisiana. Condor 97: 438–444.

35. Cuthill IC, Maddocks SA, Weall CV, Jones EKM (2000) Body mass regulation in response to changes in feeding predictability and overnight energy expenditure. Behavioral Ecology 11: 189–195.

36. Hurly TA (1992) Energetic reserves of Marsh Tits (Parus palustris) - food and fat storage in response to variable food supply. Behavioral Ecology 3: 181–188.

37. Cohen AA, Hau M, Wikelski M (2008) Stress, metabolism, and antioxidants in two wild passerine bird species. Physiological and Biochemical Zoology 81: 463–472.

38. Fokidis HB, Greiner EC, Deviche P (2008) Interspecific variation in avian blood parasites and haematology associated with urbanization in a desert habitat. Journal of Avian Biology 39: 300–310.

39. Fokidis HB, Orchinik M, Deviche P (2009) Corticosterone and corticosteroid binding globulin in birds: Relation to urbanization in a desert city. General and comparative endocrinology 160: 259–270.

40. Fokidis HB, Orchinik M, Deviche P (2011) Context-specific territorial behavior in urban birds: No evidence for involvement of testosterone or corticosterone. Hormones and behavior 59: 133–143.

41. Romero LM, Reed JM (2005) Collecting baseline corticosterone samples in the field: is under 3 min good enough? Comparative Biochemistry and Physiology a-Molecular & Integrative Physiology 140: 73–79.

42. Wingfield JC, Deviche P, Sharbaugh S, Astheimer LB, Holberton R, et al. (1994) Seasonal-changes of the adrenocortical responses to stress in redpolls, Acanthis flammea, in Alaska Journal of Experimental Zoology 270: 372–380.

43. Wingfield JC, Moore IT, Vasquez RA, Sabat P, Busch S, et al. (2008) Modulation of the adrenocortical responses to acute stress in northern and southern populations of Zonotrichia. Ornitologia Neotropical 19: 241–251.

44. Pyle P (1997) Identification guide to North American birds, Part I: Columbidae to Ploceidae: Slate Creek Press. 732 p.

45. Salewski V, Kery M, Herremans M, Liechti F, Jenni L (2009) Estimating fat and protein fuel from fat and muscle scores in passerines. Ibis 151: 640–653.

46. Bergstrom BJ, Sherry TW (2008) Estimating lipid and lean body mass in small passerine birds using TOBEC, external morphology and subcutaneous fat-scoring. Journal of Avian Biology 39: 507–513.

47. Deviche P, Breuner C, Orchinik M (2001) Testosterone, corticosterone, and photoperiod interact to regulate plasma levels of binding globulin and free steroid hormone in Dark-eyed Juncos, Junco hyemalis. General and comparative endocrinology 122: 67–77.

48. Rosner W, Hryb DJ, Khan MS, Nakhla AM, Romas NA (1991) Sex hormone-binding globulin-anatomy and physiology of a new regulatory system. Journal of Steroid Biochemistry and Molecular Biology 40: 813–820.

49. Breuner CW, Orchinik M (2002) Plasma binding proteins as mediators of corticosteroid action in vertebrates. Journal of Endocrinology 175: 99–112.

50. Orchinik M, Matthews L, Gasser PJ (2000) Distinct specificity for corticosteroid binding sites in amphibian cytosol, neuronal membranes, and plasma. General and comparative endocrinology 118: 284–301.

51. Breuner CW, Orchinik M (2001) Seasonal regulation of membrane and intracellular corticosteroid receptors in the house sparrow brain. Journal of Neuroendocrinology 13: 412–420.

52. Barsano CP, Baumann G (1989) Simple algebraic and graphic methods for the apportionment of hormone (and receptor) into bound and free fractions in binding equilibria - or how to calculate bound and free hormone. Endocrinology 124: 1101–1106.

53. Guglielmo CG, Cerasale DJ, Eldermire C (2002) A field test of plasma lipid metabolites as measures of stopover habitat quality for migratory birds. Integrative and Comparative Biology 42: 1238–1238.

54. Guglielmo CG, Cerasale DJ, Eldermire C (2005) A field validation of plasma metabolite profiling to assess refueling performance of migratory birds. Physiological and Biochemical Zoology 78: 116–125.

55. Petraitis PS, Dunham AE, Niewiarowski PH (1996) Inferring multiple causality: The limitations of path analysis. Functional Ecology 10: 421–431.

56. Myers RH (1990) Classical and modern regression with applications. Boston: PWS Kent.

57. Stavig GR (1981) Multiple comparison tests for path analysis and multiple regression. Journal of Experimental Education 50: 39–41.

58. Browne MW, Cudeck R (1992) Alternative ways of assessing model fit. Sociological Methods & Research 21: 230–258.

59. Bentler PM, Bonett DG (1980) Significance tests and goodness of fit in the analysis of covariance-structures. Psychological Bulletin 88: 588–606.

60. Holberton RL, Wilson CM, Hunter MJ, Cash WB, Sims CG (2007) The role of corticosterone in supporting migratory lipogenesis in the dark-eyed junco, Junco hyemalis: A model for central and peripheral regulation. Physiological and Biochemical Zoology 80: 125–137.

61. Meijer T, Mohring FJ, Trillmich F (1994) Annual and daily variation in body mass and fat of starlings Sturnus vulgaris. Journal of Avian Biology 25: 98–104.

62. Sharbaugh SM (2001) Seasonal acclimatization to extreme climatic conditions by black-capped chickadees (Poecile atricapilla) in interior Alaska (64 degrees N). Physiological and Biochemical Zoology 74: 568–575.

63. Balasse EO (1979) Kinetics of ketone body metabolism in fasting humans. Metabolism-Clinical and Experimental 28: 41–50.

64. Caamano GJ, Sanchezdelcastillo MA, Linares A, Garciaperegrin E (1990) In vivo lipid and amino acid synthesis from 3-hydroxybutyrate in 15 day old chick. Archives Internationales De Physiologie De Biochimie Et De Biophysique 98: 217–224.

65. Coppack SW, Jensen MD, Miles JM (1994) In vivo regulation of lipolysis in humans. Journal of Lipid Research 35: 177–193.

66. Scharrer E (1999) Control of food intake by fatty acid oxidation and ketogenesis. Nutrition 15: 704–714.

67. Breuner CW, Lynn SE, Julian GE, Cornelius JM, Heidinger BJ, et al. (2006) Plasma-binding globulins and acute stress response. Hormone and Metabolic Research 38: 260–268.

68. Malisch JL, Breuner CW (2010) Steroid-binding proteins and free steroids in birds. Molecular and Cellular Endocrinology 316: 42–52.

69. Petersen HH, Andreassen TK, Breiderhoff T, Brasen JH, Schulz H, et al. (2006) Hyporesponsiveness to glucocorticoids in mice genetically deficient for the corticosteroid binding globulin. Molecular and Cellular Biology 26: 7236–7245.

70. Malisch JL, Kelly SA, Bhanvadia A, Blank KM, Marsik RL, et al. (2009) Lines of mice with chronically elevated baseline corticosterone levels are more susceptible to a parasitic nematode infection. Zoology 112: 316–324.

71. Love OP, Breuner CW, Vezina F, Williams TD (2004) Mediation of a corticosterone-induced reproductive conflict. Hormones and behavior 46: 59–65.

72. Dallman M, Pecoraro N, Warne J, Akana S (2009) Stress, Glucocorticoids & Insulin: Scaffold for Obesity. Biological Psychiatry 65: 176S–176S.

73. Watts LM, Manchem VP, Leedom TA, Rivard AL, McKay RA, et al. (2005) Reduction of hepatic and adipose tissue glucocorticoid receptor expression with antisense oligonucleotides improves hyperglycemia and hyperlipidemia in diabetic rodents without causing systemic glucocorticoid antagonism. Diabetes 54: 1846–1853.

74. Scanes CG (2009) Perspectives on the endocrinology of poultry growth and metabolism. General and comparative endocrinology 163: 24–32.

75. Macfarlane DP, Forbes S, Walker BR (2008) Glucocorticoids and fatty acid metabolism in humans: fuelling fat redistribution in the metabolic syndrome. Journal of Endocrinology 197: 189–204.

76. Thurston RJ, Bryant CC, Korn N (1993) The effects of corticosterone and catecholamine infusion on plasma-glucose levels in chicken (Gallus domesticus) and turkey (Meleagris gallapavo). Comparative Biochemistry and Physiology C-Pharmacology Toxicology & Endocrinology 106: 59–62.

77. Cyr NE, Earle K, Tam C, Romero LM (2007) The effect of chronic psychological stress on corticosterone, plasma metabolites, and immune responsiveness in European starlings. General and comparative endocrinology 154: 59–66.

78. Remage-Healey L, Romero LM (2001) Corticosterone and insulin interact to regulate glucose and triglyceride levels during stress in a bird. American Journal of Physiology-Regulatory Integrative and Comparative Physiology 281: R994–R1003.

79. Remage-Healey L, Romero LM (2002) Corticosterone and insulin interact to regulate plasma glucose but not lipid concentrations in molting starlings. General and comparative endocrinology 129: 88–94.

80. Swain SD (1991) Metabolism and energy reserves of brood-rearing horned larks (Eremophila alpestris). Comparative Biochemistry and Physiology a-Physiology 99: 69–73.

81. Jenni-Eiermann S, Jenni L, Kvist A, Lindstrom A, Piersma T, et al. (2002) Fuel use and metabolic response to endurance exercise: a wind tunnel study of a long-distance migrant shorebird. Journal of Experimental Biology 205: 2453–2460.

82. Lindgard K, Stokkan KA, Lemaho Y, Groscolas R (1992) Protein-utilization during starvation in fat and lean svalbard ptarmigan (Lagopus-mutus-hyperboreus). Journal of Comparative Physiology B-Biochemical Systemic and Environmental Physiology 162: 607–613.

83. Jenni-Eiermann S, Jenni L (1994) Plasma metabolite levels predict individual body-mass changes in a small long-distance migrant, the garden-warbler. Auk 111: 888–899.

84. Klandorf H, Probert IL, Iqbal M (1999) In the defence against hyperglycaemia: an avian strategy. Worlds Poultry Science Journal 55: 251–268.

85. Tsahar E, Arad Z, Izhaki I, Guglielmo CG (2006) The relationship between uric acid and its oxidative product allantoin: a potential indicator for the evaluation of oxidative stress in birds. Journal of Comparative Physiology B-Biochemical Systemic and Environmental Physiology 176: 653–661.

86. Gate L, Paul J, Ba GN, Tew KD, Tapiero H (1999) Oxidative stress induced in pathologies: the role of antioxidants. Biomedicine & Pharmacotherapy 53: 169–180.

87. Buffenstein R, Edrey YH, Yang T, Mele J (2008) The oxidative stress theory of aging: embattled or invincible? Insights from non-traditional model organisms. Age 30: 99–109.

88. Cohen A, Klasing K, Ricklefs R (2007) Measuring circulating antioxidants in wild birds. Comparative Biochemistry and Physiology B-Biochemistry & Molecular Biology 147: 110–121.

89. Smith CL, Toomey M, Walker BR, Braun EJ, Wolf BO, et al. (2011) Naturally high plasma glucose levels in mourning doves (Zenaida macroura) do not lead to high levels of reactive oxygen species in the vasculature. Zoology 114: 171–176.

90. Ku HH, Sohal RS (1993) Comparison of mitochondrial pro-oxidant generation and antioxidant defenses between rat and pigeon-possible basis of variation in longevity and metabolic potential. Mechanisms of Ageing and Development 72: 67–76.

